# Quantification of Cholesterol Incorporation in Giant Unilamellar Vesicles Produced by a Modified cDICE Method

**DOI:** 10.1101/2025.11.10.687550

**Authors:** Marcos Arribas Perez, Gijsje H. Koenderink

## Abstract

Cholesterol is an essential component of eukaryotic cell membranes, influencing membrane packing, fluidity, and domain formation. Replicating these properties in model membranes is critical for reconstitution studies, but common emulsion-based methods for producing giant unilamellar vesicles (GUVs) fail to incorporate cholesterol efficiently. Here, we use methyl-β- cyclodextrin–cholesterol (MβCD-CL) complexes to deliver cholesterol into GUVs produced by the emulsion droplet interface crossing encapsulation (eDICE) method and demonstrate a convenient way to quantify the degree of cholesterol incorporation using fluorescent membrane biosensors. Spectral imaging of NR12A as well as fluorescence lifetime imaging of Flipper-TR revealed dose-dependent increases in cholesterol content for DOPC GUVs upon MβCD-CL addition, consistent with increased membrane order. By calibrating these effects against GUVs with defined cholesterol contents prepared via gel-assisted swelling, we found that the cholesterol content of eDICE vesicles can be increased to at least 40 mol%. Binary mixtures of DOPC with saturated lipids (DMPC and PC(18:0-14:0)) showed a similar trend as pure DOPC GUVs. Interestingly, we could trigger liquid-ordered domain formation by adding cholesterol to DOPC:DMPC vesicles. Our findings provide a quantitative and non-disruptive method to modulate and assess cholesterol content in emulsion-based GUVs, advancing their use in bottom-up synthetic biology and membrane biophysics.

## 1. Introduction

Cell membranes are dynamic structures that, far from being simple barriers, play key roles in a wide range of cellular activities, including compartmentalization, signal transduction, membrane trafficking, and cell shape regulation ^1^. They are composed of a diverse mixture of lipids, proteins, and glycolipids, organized in a highly regulated and heterogeneous manner ^1, 2^. The lipid bilayer itself contains hundreds of distinct lipid species, which are asymmetrically distributed across the two leaflets and contribute to the structural and functional complexity of cell membranes ^3^. A key component of eukaryotic plasma membranes is cholesterol, which constitutes between 10 and 30% of the total lipid content ^1^. Cholesterol critically influences membrane characteristics including thickness, rigidity, packing, and permeability ^4^.

To study the biophysical properties of cell membranes in a controlled manner, giant unilamellar vesicles (GUVs) have become a widely used biomimetic model system. These cell-sized lipid vesicles mimic the architecture of biological membranes and allow control over the membrane physicochemical properties as well as the reconstitution of minimal cellular processes under controlled environments ^5, 6^. Achieving cholesterol incorporation is essential for generating membrane models that faithfully replicate the composition and physical properties of cellular membranes. Furthermore, in GUVs made of mixtures with saturated and unsaturated lipids, cholesterol supports the formation of coexisting liquid- ordered (Lo) and liquid-disordered (Ld) domains ^7^. These phase-separated membranes can serve as tool to spatially organize membrane-associated proteins in bottom-up reconstitution studies. For instance, protein condensates and actin networks have been shown to preferentially associate with either Lo or Ld phases ^8, 9^.

However, the efficiency of cholesterol incorporation in GUVs is highly dependent on the preparation technique. Using mass spectrometry, a recent study has shown that swelling-based vesicle preparation methods showed a robust and reproducible integration of cholesterol into the bilayer with deviations of less than 5 mol% from the stock lipid mixture. In contrast, the GUVs made by the emulsion-transfer method contained less than 20% of the cholesterol expected from the lipid stocks ^10^. Even poorer cholesterol incorporation (<10%) has been reported for cDICE ^11^, another emulsion-based method. A modified version of the cDICE method (double layer cDICE), where an additional silicone oil layer (free of mineral oil) with the cholesterol dissolved was added to the standard cDICE protocol, was reported to increase the incorporation of cholesterol into GUVs, but still only to about 25-30% of the total amount added to the lipid mixture ^12^. Emulsion-based methods for vesicle fabrication have important advantages over swelling-based methods because they allow for accurate encapsulation of cytosolic contents, and are therefore popular for complex reconstitutions such as efforts to build synthetic cell ^13^. Unfortunately, the poor cholesterol incorporation limits the ability of emulsion-based vesicles to reproduce the biophysical characteristics of native cell membranes.

While mass spectrometry allows the precise quantification of different lipid species in the membrane, it has the drawback of being destructive. Moreover, it provides an ensemble-averaged readout and cannot be used to test for heterogeneities in vesicle populations. An interesting alternative probe of lipid composition is provided by environment-sensitive fluorescent probes that respond to changes in lipid packing and lateral pressure by altering their fluorescence lifetime or spectral profile. Solvatochromic probes, such as Laurdan, Pro12A, di-4-ANEPPDHQ and Nile Red derivatives (NR12S, NR12A and NR4A) have been extensively used to quantify lipid packing within lipid membranes ^14–19^. These probes show a spectral shift in response to the polarity of their local environment. In lipid membranes the polarity represents the hydration level, which is directly related to the membrane order. Saturated lipids pack tightly and leave less space for water molecules at the interface between the hydrophobic and hydrophilic regions of the membrane while unsaturated lipids pack more loosely, creating more disordered membranes that allow more water to enter this region ^20^. A different but complementary environment-sensitive probes is the planarizable push-pull probe Flipper-TR. Contrary to the solvatochromic dyes that respond to polarity, Flipper-TR is a mechanosensitive probe that changes conformation in response to lateral pressure in the bilayer. Its transition from a twisted to a planar conformation is accompanied by an increase in its fluorescence lifetime ^21^. Flipper-TR has been used to investigate lipid packing and phase behaviour in model membranes and as a membrane tension reporter in living cells ^17, 22^.

In this study, we use a recent modification of the cDICE method (eDICE ^23^) to create GUVs, followed by the delivery of cholesterol *after* vesicle formation using methyl-β- cyclodextrin–cholesterol complexes, a method previously shown to enable cholesterol insertion into membranes ^11^. To assess the extent of cholesterol incorporation, we employ a combination of *in situ* fluorescence microscopy techniques using environment-sensitive probes that report on lipid packing and membrane order. This approach provides a quantitative and minimally invasive method to quantify the extent of cholesterol integration in the membranes with the final aim to achieve a more accurate representation of biological membranes in vesicle-based model systems.

## 2. Materials and methods

### 2.1. Materials

Mineral oil (M5904), chloroform (372978), biotinylated bovine serum albumin (A8549), Poly-vinyl-alcohol (PVA) 145 kDa (8.14894) Sigma-Aldrich, and Methyl-β-cyclodextrin (MβCD) (C4555) were purchased from Sigma. Decane (10497480) and Neutravidin (10443985) were bought from Thermo Fisher Scientific. Silicone oil 5cSt (7844) was ordered from Carl Roth. Lipids, specifically cholesterol (ovine) (700000P), 1,2-dimyristoyl-sn-glycero-3-phosphocholine (DMPC, 14:0 PC) (850345), 1,2-dioleoyl-sn-glycero-3- phosphocholine (DOPC) (850375), 1-stearoyl-2-myristoyl-sn-glycero-3-phosphocholine (PC(18:0-14:0)) (850464), and 1,2-dioleoyl-sn-glycero-3-phosphoethanolamine-N- (biotinyl) (sodium salt) (Biotin-PE), were purchased from Avanti Polar Lipids. Flipper-TR (SC020) from Spirochrome AG and MemGlow NR12A (MG07) from Cytoskeleton, Inc. were supplied by Tebu Bio.

### 2.2. Gel assisted swelling

To make GUVs using the gel swelling method we followed a protocol described before ^24^. We prepared a 5% (w/v) solution of poly-vinyl alcohol (PVA) in a solution of 200mM sucrose in milliQ water. We coated a 24x24mm glass coverslip with 100 µL of the PVA solution and removed the excess by tilting the glass on a tissue to minimize gel thickness. The PVA-coated coverslip was heated at 50°C for 30 minutes in an oven to dry the gel. Then, 10 µL of the lipid mixture (with different compositions, see below) dissolved in chloroform at a total lipid concentration of 1.25 mM were spread on the dried PVA. The organic solvent was evaporated by placing the coverslips under vacuum for 30 minutes. Next, 300 µL of a swelling buffer containing 15mM Tris pH 7.4 and 140 mM sucrose (175 mOsm/kg) were carefully added on top of the gel. After 1 hour incubation at room temperature, the GUVs were collected and diluted 6 times in observation buffer (15 mM Tris pH 7.4 and 170 mM glucose, 200 mOsm/kg). The osmolarity of the buffers was measured with a freezing point osmometer (Osmomat 3000, Gonotec, Germany). One set of lipid mixtures used for the gel swollen GUVs was based on DOPC:cholesterol in molar ratios of 100:0, 90:10, 75:25 and 60:40. In addition we also prepared samples with binary lipid mixtures of DOPC:DMPC (60:40) and DOPC: PC(18:0-14:0) (60:40), and ternary lipid mixtures of DOPC:DMPC:cholesterol (36: 24: 40) and DOPC:PC(18:0- 14:0):cholesterol (36: 24: 40). The GUVs were used for confocal imaging within 3 hours after fabrication.

### 2.3. eDICE

To produce GUVs using emulsion droplet interface crossing encapsulation (eDICE) we followed a previously described method ^23^. eDICE is an adaptation of the original cDICE method ^25^ where, instead of using a capillary to produce water-in-oil emulsion droplets, we use mechanical agitation to create the droplets. Briefly, the desired lipids were mixed in chloroform and dried under a N2 stream. We used three different lipid mixes: DOPC:Biotin-PE (99:1 mol%), DOPC: PC(18:0-14:0):Biotin-PE (59:40:1 mol%) and DOPC:DMPC:Biotin-PE (59:40:1 mol%). The amount of lipids dried (1.75 µmol) yields a final lipid concentration in the lipid-in-oil-mix of 0.25 mM. The vials with the dried lipids were transferred into a glovebox to keep the humidity of the environment below 1%, as this has been reported to improve the quality of the membranes and the vesicle yield ^26^. Once in the glovebox, the lipids were resuspended in 50 µL of chloroform, 400 µL of decane and 6.5 mL of a mix of silicone oil and mineral oil (8:2 vol/vol). The vials were sealed and taken out of the glove box, and the lipid-in-oil mixtures were sonicated for 15 minutes on ice in a bath sonicator. The GUVs were prepared in a room where the humidity was kept below 40% using a custom-made spinning table and a 3D-printed chamber described in a previous publication^26^ . We placed the chamber on the spinning table and made it rotate at 2000 rpm. We added 700 µL of an outer aqueous solution (OAS) containing 190 mM glucose (200 mOsm/kg) into the spinning chamber followed by 5 mL of the lipid-in-oil mix. Then, 1 mL of the lipid-in-oil mix was set aside into a 2ml Eppendorf tube. To create the droplet emulsion we added 25 µL of the inner aqueous solution (IAS) to the tube, with a composition chosen to be compatible with protein encapsulation for future studies (20 mM Tris-HCl pH 7.4, 50 mM KCl, 2 mM MgCl2 and 6.5 % vol/vol Optiprep, 175 mOsm/kg). We manually created the emulsion by scratching the tube over a tube holder 10-15 times. The emulsion was pipetted into the spinning chamber and centrifuged for 3 minutes. Once stopped, we carefully pipetted off the excess of oil from the chamber and added 250 µl of 60 mM Tris-HCl pH 7.4 and 100 mM glucose solution isosmotic with the OAS (200 mOsm/Kg) to stabilize the pH of the outer solution. The chambers were kept tilted at an angle of approximately 45° for 10 minutes to allow the GUVs to sediment to the bottom. The GUVs were then collected from the bottom of the spinning chamber with a 200µL pipette with a cut-off tip to minimize shear forces and transferred to a fresh Eppendorf tube. Finally, the GUVs were diluted by a factor of 4 in observation buffer (15 mM Tris-HCl pH 7.4 and 170 mM glucose). The GUVs were used for confocal imaging experiments within 3 hours after fabrication.

### 2.4. GUV labelling

The fluorescent dyes were dissolved in dimethylsulfoxide (DMSO) following the manufacturer’s instructions (Flipper-TR stock at 1mM and NR12A at 20 µM) and stored in small aliquots at -20° C. In all the experiments presented here, the GUVs were labelled after being formed by incubating them for 15 minutes with the fluorescent dye of interest. The concentration of dye used to label the membrane of the GUVs was 1 µM for Flipper-TR (following the manufacturer’s recommendation) and 0.1 µM for NR12A (in line with previous reports^17, 27^).

### 2.5. Cholesterol delivery to the membrane

Methyl-β-cyclodextrin (MβCD) was preloaded with cholesterol following a previously described protocol ^11^. We first dissolved 2 mg of cholesterol and 55 mg of MβCD in 500 µl of methanol:chloroform (4:1 vol/vol) in a 8mL glass vial and then dried the mixture under nitrogen followed by vacuum for one hour. This results in the formation of crystals at the bottom of the tube. The crystals were resuspended in 2 mL of observation buffer heated at 80°C by vortexing vigorously until they were completely resuspended, resulting in a suspension of MβCD-cholesterol complexes at a concentration of 2.6 mM cholesterol (1mg/ml) and 8.3 mM MβCD (11 mg/ml). The suspension was stored at -20°C and used for no longer than two weeks.

The GUVs were incubated with the MβCD-cholesterol complexes for 15 minutes before imaging. For clarity, we refer to the concentration of cholesterol added to the GUVs, but we emphasize that the cholesterol is always added complexed with MβCD. In our experiments we incubated the GUVs with 10 µM cholesterol (32 µM MβCD), 30 µM cholesterol (96 µM MβCD) and 100 µM cholesterol (320 µM MβCD).

### 2.6. Microscopy

All confocal spectral imaging and fluorescence lifetime microscopy measurements were performed on an inverted Leica STELLARIS 8 FALCON laser scanning confocal microscope equipped with a white light laser and a 63x glycerol immersion objective (HC PL APO, WD0.3 mm, NA 1.2). GUVs were observed in Ibidi 18-well µ-slides (Ibidi catalog # 81817). To prevent rupture of the GUVs and reduce drift while imaging, the slides were treated with a solution of biotinylated bovine serum albumin (BSA-biotin, 1 mg/mL in milliQ water) for 15 minutes, followed by washing with observation buffer and another 10 min incubation with a neutravidin solution (1 mg/mL in observation buffer). Afterwards, the slide was washed twice with observation buffer to remove unbound neutravidin. The wells in the microscope slide were left filled with 50 µL of observation buffer to prevent drying of the surface. The GUVs (50 µL) were added directly to the observation buffer in the well and allowed to sink down for 5 minutes before imaging.

### 2.7. Confocal Spectral imaging and GP analysis of Nile Red 12A

To quantify the generalized polarization (GP) parameter of the NR12A probe within the membrane of the GUVs we performed confocal spectral imaging ^14, 28^. The GUVs labelled with NR12A were excited at 540 nm (laser power 15%) and the emission was acquired between 560 and 700 nm with a spectral step of 10 nm per channel using the lambda scan function of the microscope. The emission was collected using a HyD detector set to counting mode (internal gain 2.5%). The scan speed was set to 700 Hz and the image resolution at 256 x 256 pixels.

The Generalized Polarization (GP) was computed from the fluorescence spectral images using a custom Python script based on standard libraries, including NumPy, scikit-image, pandas, matplotlib, and tifffile. Prior to GP analysis, the fluorescence spectral images were pre-processed using the StackReg plugin of Fiji to ensure that the GUVs are in the same position in every channel of the image ^29^. The images were loaded as tiff stacks and each stack was converted to a 2D overlay image using maximum intensity projection to create a mask. High-intensity outliers (e.g., saturated pixels) were excluded by applying a brightness threshold. The resulting image was contrast-adjusted based on user-defined percentiles and segmented using automatic thresholding via the Otsu algorithm to generate a binary mask. This mask was used to isolate relevant regions for GP analysis, excluding bright artifacts.

Pixel-wise fluorescence intensities were extracted from the selected slices of the stack corresponding to the two wavelengths of interest (570 nm and 640 nm). The intensities at the selected wavelengths were used to calculate the pixel-wise generalised polarization (GP) according to the following equation^14^:

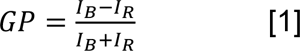

where *I_B_* is the fluorescence intensity at the shorter wavelength and *I_R_* is the fluorescence intensity at the longer wavelength. The GP values were mapped back onto the original image geometry to generate GP images. From the pixel-wise GP values we calculated the mean GP of the vesicle in the image.

### 2.8. Fluorescence lifetime microscopy and analysis for Flipper-TR

For FLIM measurements of Flipper-TR, samples were excited at 488 nm using 50% laser power. Emission was collected in the 575–625 nm range using a HyD detector in photon-counting mode (internal gain 10%). The laser power was adjusted to ensure a photon count rate below 0.5 photons per excitation pulse to minimise pile-up artifacts where more than 1 photon per pulse are detected. To ensure the acquisition of enough photons per pixel to obtain reliable decay fits without affecting the photon rate per pulse, we used 5 frame repetitions per image. All acquisitions were performed at a laser repetition rate of 20 MHz.

Fluorescence lifetime analysis was conducted using the Leica Application Suite LAS X FLIM/FCS software (version 4.5.0). Fluorescence decay curves were fitted using a double-exponential reconvolution model, employing the instrument response function (IRF) generated by the FALCON FLIM module within a fitting window of 0.2–45 ns ^17^. Reported lifetime values correspond to the mean intensity-weighted lifetime (*τ_m int_*), calculated by the software according to the following equation^17^:

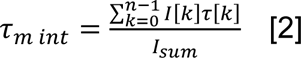

Where *I[k]* is the intensity of each exponential component, *τ[k]* the corresponding lifetime, and *I_sum_* the total fluorescence intensity.

Phasor analysis was also carried out using the LAS X FLIM/FCS software. A wavelet filter with a threshold of 50 was applied to enhance the distinction of photon clouds in the phasor plot. For quantification, the centre of each photon cloud was manually selected using the circular selection tool with a radius of 20.

### 2.9. Statistics and calibration curves

Spectral and lifetime data for eDICE GUVs with different amounts of MβCD–CL added in the outer medium were based on at least 50 GUVs from 3 different replicates per lipid composition. Prior to statistical comparison, the data were evaluated for normality and homogeneity of variance. As these conditions were not met, we used a non-parametric Kruskal–Wallis ANOVA test to evaluate the statistical significance between the samples incubated with different concentrations of MβCD–CL.

For the calibration analysis, spectral and lifetime measurements were acquired from gel- swollen GUVs prepared with defined cholesterol concentrations. Each condition included at least 30 GUVs from 3 different replicates per lipid composition. The resulting lifetime (Flipper-TR) or GP values (y) as a function of MβCD–CL concentration added in the outer medium (x)were fitted by a linear regression (*y=c+mx*) and the fitting parameters *c* and *m* were subsequently used to estimate the effective cholesterol content in the eDICE GUVs following incubation with varying concentrations of MβCD–CL.

All statistical analyses and curve fittings were performed using OriginPro (OriginLab, Northampton, MA, USA).

## 3. Results and discussion

### 3.1. Cholesterol delivery by MβCD into DOPC GUVs made by eDICE can be sensed by NR12A and Flipper-TR probes

We first investigated the cholesterol incorporation in GUVs made of a single lipid species (DOPC) that forms a homogeneous fluid membrane. To deliver cholesterol into the membrane of GUVs prepared by the eDICE method we incubated the vesicles with different concentrations of MβCD-CL complexes following a previous report ^11^. For clarity we will indicate the concentration of cholesterol in the sample when we added the MβCD- CL complexes. The degree of cholesterol incorporation into the membranes was assessed *in situ* by measuring changes in the lipid packing and lateral pressure through two complementary environment-sensitive fluorescence probes.

The incorporation of cholesterol in the membrane of the eDICE-produced GUVs upon incubation with MβCD-CL complexes was first evaluated by looking at changes in the generalised polarization (GP) of the polarity sensitive probe NR12A using spectral confocal imaging. The incorporation of cholesterol in the DOPC membrane is expected to increase the lipid packing ^30, 31^, leading to a more hydrophobic environment around the NR12A molecules that should result in a spectral shift of the fluorescence emission of the dye towards lower wavelengths^16^ (Figure 1a). Our spectral imaging measurements using NR12A indeed reveal a clear, concentration-dependent increase in the generalized polarization (GP) values of DOPC GUVs upon the addition of MβCD-CL complexes to the outer medium (Figure 1b-c). The GP images show different GP values along the membrane of the GUVs (Figure 1b). These differences are not due to inhomogeneous membrane order but result from an imaging artifact known as photoselection ^28, 32^. The excitation of the probe with a linearly polarized light leads to a brighter fluorescence emission of NR12A in regions aligned in parallel with the light polarization and dimmer fluorescence in perpendicular regions. These brightness variations translate into artificial GP inhomogeneities. Importantly, the same effect is consistently observed in all GUVs, independent of the sample condition. GP values were averaged along the vesicle contour, which is expected to smooth out orientation-dependent variations while preserving the relative changes between conditions. The averaged GP value of -0.51 ± 0.05 for pure DOPC GUVs reflects a highly disordered membrane environment, typical of unsaturated lipid bilayers ^33^. Upon exposure to 10 µM MβCD-CL, the GUVs show a subtle increase of the mean GP (reaching -0.44 ± 0.08, implying a shift of ΔGP=0.07). Further increasing the MβCD-CL concentration results in a further increase of the NR12A GP, reaching mean GP values of -0.25 ± 0.09 (ΔGP=0.26) and -0.10 ± 0.11 (ΔGP=0.41) after incubation with 30 µM and 100 µM cholesterol, respectively (Figure 1c). This concentration-dependent rise in the GP value clearly suggests that cholesterol has been delivered to the membranes. Compared to the GUVs not exposed to cholesterol, the distribution of the GP values in the samples incubated with MβCD-CL is broader and shows some outliers indicating that the cholesterol incorporation level varies across the GUVs in the sample.

**Figure 1.**
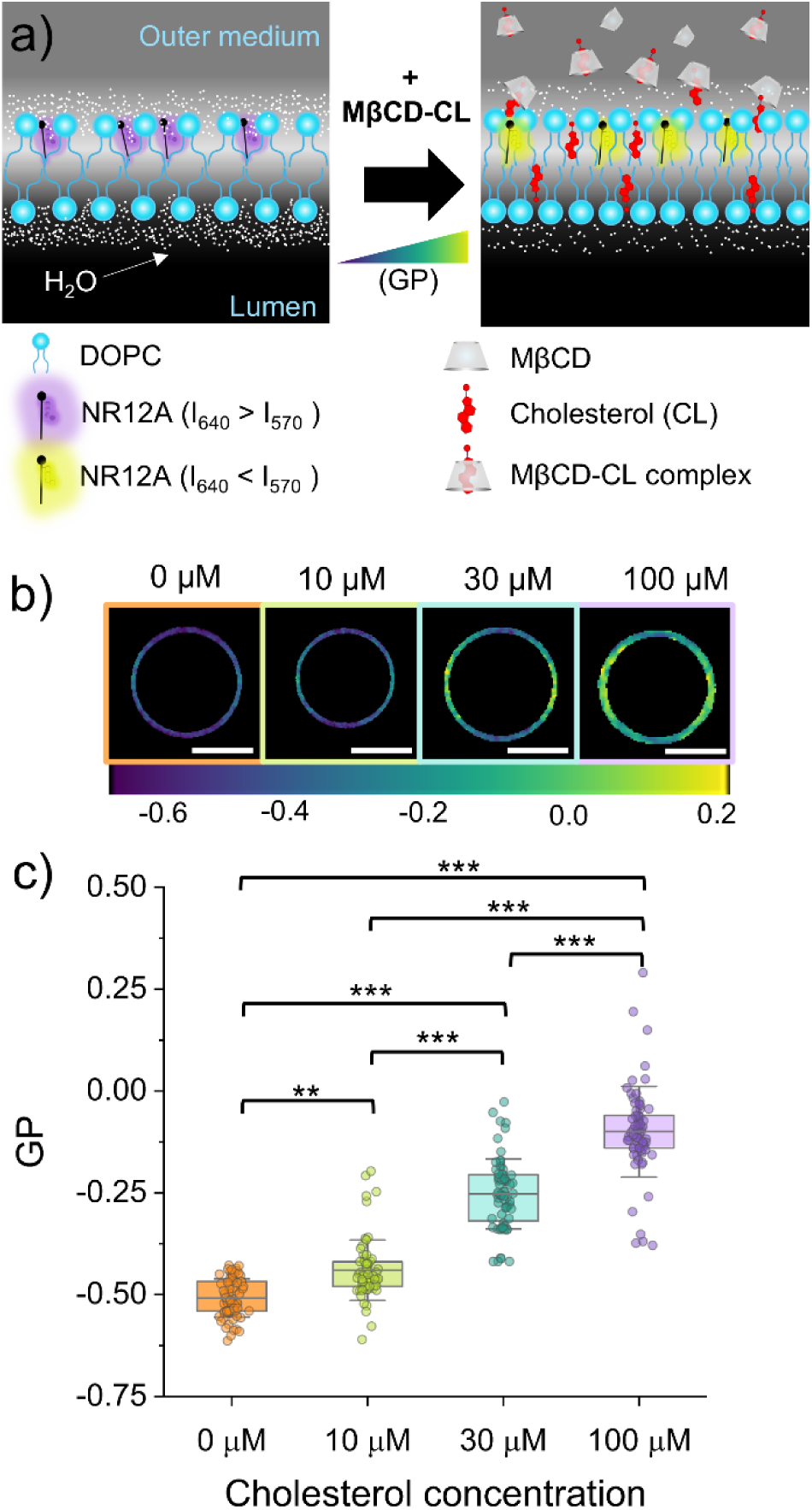
NR12A generalized polarization (GP) as a reporter of cholesterol incorporation in DOPC GUVs. a) Schematic representation of the change in GP of NR12A in DOPC membranes upon MβCD-driven delivery of cholesterol in the outer medium. b) Representative color-coded GP images generated from confocal images at the equatorial plane of eDICE GUVs at different cholesterol (MβCD-CL) concentrations. Scale bars indicate 10µm. c) GP measured in eDICE GUVs at the different conditions. Box percentile 5-95, line in box is the mean and whiskers the SD. Points are individual GUVs (at least 50 per condition from at least 3 different replicates). Statistical significance was tested using a non-parametric Kruskal-Wallis test (** p< 0.01 and *** p< 0.001).

To independently verify our observations, we used a complementary approach based on fluorescence lifetime microscopy (FLIM) in combination with the Flipper-TR probe to detect the addition of cholesterol into the membranes. Unlike the NR12A probe that responds to polarity, Flipper-TR responds to changes in membrane packing ^21^. The incorporation of cholesterol in the DOPC membrane is expected to increase the lipid packing^30, 31^ and thereby increase the Flipper-TR fluorescence lifetime (Figure 2a). In all conditions the fluorescence lifetime was homogeneous along the membrane of the GUVs (Figure 2b). The fluorescence lifetime of Flipper-TR in DOPC GUVs increased progressively with higher concentrations of MβCD-CL, from 2.96 ± 0.06 ns in untreated vesicles to 3.41 ± 0.05 ns, 4.05 ± 0.13 ns, and 4.37 ± 0.20 ns after incubation with 10, 30, and 100 µM cholesterol, respectively (Figure 2a-b).

**Figure 2.**
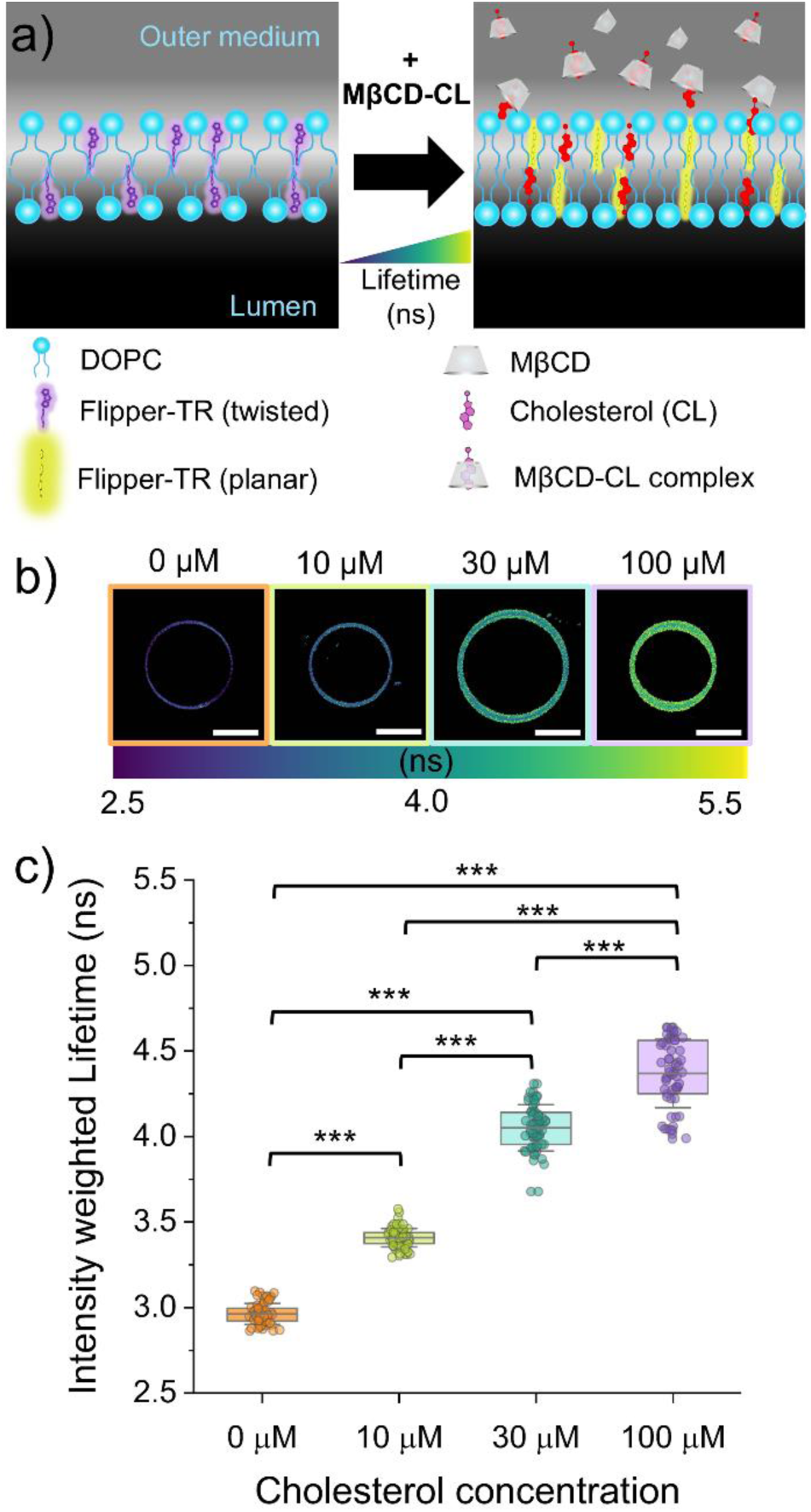
Flipper-TR fluorescence lifetimes as a reporter of cholesterol incorporation in DOPC GUVs. a) Schematic representation of the conformational change resulting in longer fluorescence lifetimes of Flipper-TR in DOPC membranes upon MβCD-driven delivery of cholesterol in the outer medium. b) Representative color-coded mean intensity weighted lifetime images at the equatorial plane of eDICE GUVs at different cholesterol (MβCD-CL) concentrations. Scale bars indicate 10 µm. c) Mean intensity weighted lifetimes of eDICE GUVs as a function of the MβCD-CL concentration in the outer medium. Box percentile 5-95, line in box is the mean and whiskers the SD. Points are individual GUVs (at least 50 per condition from at least 3 different replicates). Statistical significance was tested using a non-parametric Kruskal-Wallis test (*** p< 0.001).

To estimate the cholesterol content in eDICE GUVs after incubation with varying concentrations of MβCD-CL complexes, we prepared control GUVs by gel swelling made of DOPC combined with defined proportions of cholesterol (0%, 10%, 25%, and 40%). Swelling methods are known to enable membrane formation from diverse lipid mixtures, including synthetic, charged, and natural lipids, as well as high cholesterol contents^13^. For instance, swelling assisted by an electric field (electroformation) has previously been shown to give complete incorporation with a deviation from the stock solution lower than 5%^10^. The corresponding GP values obtained for these samples were -0.53 ± 0.04, -0.45 ±, -0.30 ± 0.07, and -0.09 ± 0.09, respectively (Figure 3a). Similar changes in the GP of NR12A have been reported before in electroformed GUVs made of POPC and increasing proportions of cholesterol ^27^. In that study they observed that, compared to pure POPC vesicles, the presence of 10% of cholesterol in POPC GUVs results in a mild spectral shift of NR12A (ΔGP= 0.06) whereas GUVs containing 50% cholesterol exhibited a substantially increased GP value (ΔGP= 0.54) ^27^. To obtain a reference to interpolate the cholesterol levels in the eDICE GUVs based on their measured GP values, we fitted the data for the electroformed GUVs to a linear regression (Figure 3a). In this way, we calculated that incubation with 10 µM cholesterol corresponds to a final cholesterol content of the eDICE GUVs of ∼10%, 30 µM to ∼28% and 100 µM to ∼42% cholesterol (Figure 3c).

**Figure 3.**
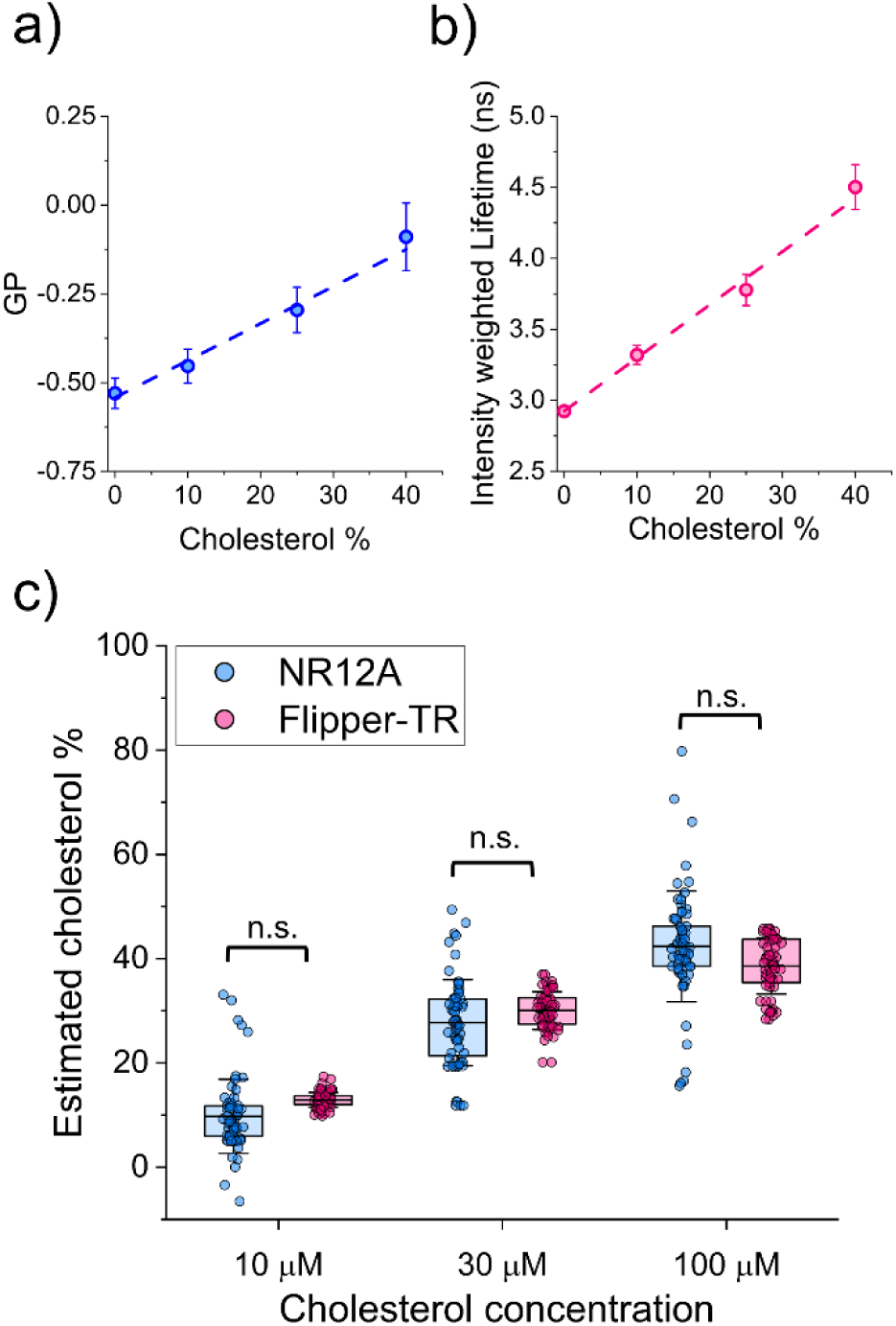
Calibration of fluorescence lifetime and GP measurements of the cholesterol content of GUVs. a) Mean GP of DOPC GUVs containing different proportions of cholesterol prepared by gel swelling. The dashed line represents the linear regression fit (y = -0.542 + 0.010x). b) Mean intensity weighted lifetimes of DOPC GUVs and different proportions of cholesterol prepared by gel swelling. The dashed line represents the linear regression fit (y = 2.925 + 0.037x). In both panels a and b, dots and whiskers represent the mean ± SD (with at least 30 GUVs from 3 different replicates measured per lipid composition). c) Cholesterol percentage in the membrane of eDICE GUVs as a function of the MβCD-CL concentration in the outer medium estimated from the calibration curves from gel swollen vesicles, measured by NR12A spectral imaging (based on the data in Figure 1) and Flipper-TR fluorescence lifetime (based on the data in Figure 2). Statistical analysis (non-parametric Kruskal-Wallis test) confirms that there is no significant difference (n.s.) in the results obtained from each method.

In a similar way as with NR12A, we again used GUVs made by gel swelling as a calibration curve for the fluorescence lifetime of Flipper-TR (Figure 3b). From that, we estimated that the GUVs incubated with 10, 30 and 100 µM cholesterol complexed with MβCD contained ∼13%, 30% and 39% cholesterol (Figure 3c). These results closely match those obtained with NR12A, supporting consistent dose-dependent cholesterol incorporation in the eDICE GUVs controlled by the concentration of MβCD-CL added to the outer medium.

### 3.2. Cholesterol delivery by MβCD also works efficiently for GUVs made of binary lipid mixtures

To test whether cholesterol delivery to eDICE GUVs also extends to more complex lipid compositions, we also tested cholesterol incorporation in GUVs containing saturated phosphatidylcholine species by preparing binary mixtures of DOPC with either DMPC or PC(18:0-14:0) at a 6:4 mol/mol ratio. DMPC and PC(18:0-14:0) differ in the length and symmetry of the acyl chains and consequently their melting temperature. DMPC contains two symmetric myristoyl (14:0) chains and has a melting temperature of ∼24 °C, while PC(18:0-14:0) has one longer stearoyl (18:0) and one myristoyl (14:0) chain, resulting in an asymmetric structure and a higher melting temperature of ∼35 °C ^34^.

Prior to cholesterol addition, the NR12A GP values were −0.38 ± 0.07 for DOPC:DMPC and −0.41 ± 0.05 for DOPC:PC(18:0–14:0) (Figure 4 a-d), both notably higher than those observed in pure DOPC membranes (-0.51 ± 0.05, Figure 1 c). This suggests that the incorporation of saturated lipids increases membrane packing even in the absence of sterols. This effect of the acyl chain saturation on the GP of NR12A has been observed before even in POPC membranes, where only one of the acyl chains of the lipids is saturated ^27^. In contrast, Flipper-TR–labelled GUVs composed of the same binary mixtures exhibited average fluorescence lifetimes (3.05 ± 0.10 ns for DOPC:DMPC and 2.98 ± 0.11 ns for DOPC:PC(18:0–14:0)) comparable to that of pure DOPC (Figure 5 a-d). Consistent with these findings, similar NR12A GP values (-0.37± 0.06 for DOPC:DMPC and -0.39± 0.07 for DOPC:PC(18:0–14:0)) and Flipper-TR fluorescence lifetimes (3.07 ± 0.08 ns for DOPC:DMPC and 3.01 ± 0.06 ns for DOPC:PC(18:0–14:0)) were observed in gel-swollen GUVs of equivalent lipid composition (Figure S1). This discrepancy likely stems from the differing sensing mechanisms of the two probes: while NR12A detects changes in local polarity linked to packing^16, 27^, Flipper-TR responds to lateral pressure changes through a conformational twist ^21, 22^. These observations imply that, although saturated lipids promote tighter packing, this effect may not generate sufficient lateral stress to trigger the mechanical response required for a change in Flipper-TR lifetime. Notably, no signs of lateral heterogeneity or phase separation were observed in either system under these conditions. This distinction between NR12A and Flipper-TR behaviour is further supported by previous studies comparing environment sensitive probes. Amaro *et al*.^18^ showed that Laurdan is an accurate and sensitive indicator of lipid order while di-4-ANEPPDHQ is sensitive to the membrane potential and its GP does not correlate with lipid packing but is specifically influenced by the presence of cholesterol. Other previous studies have reported that while both NR12A and Flipper-TR detect cholesterol content, only NR12A exhibits strong sensitivity to lipid saturation whereas Flipper-TR is relatively insensitive to acyl chain ordering ^17, 27^.

**Figure 4.**
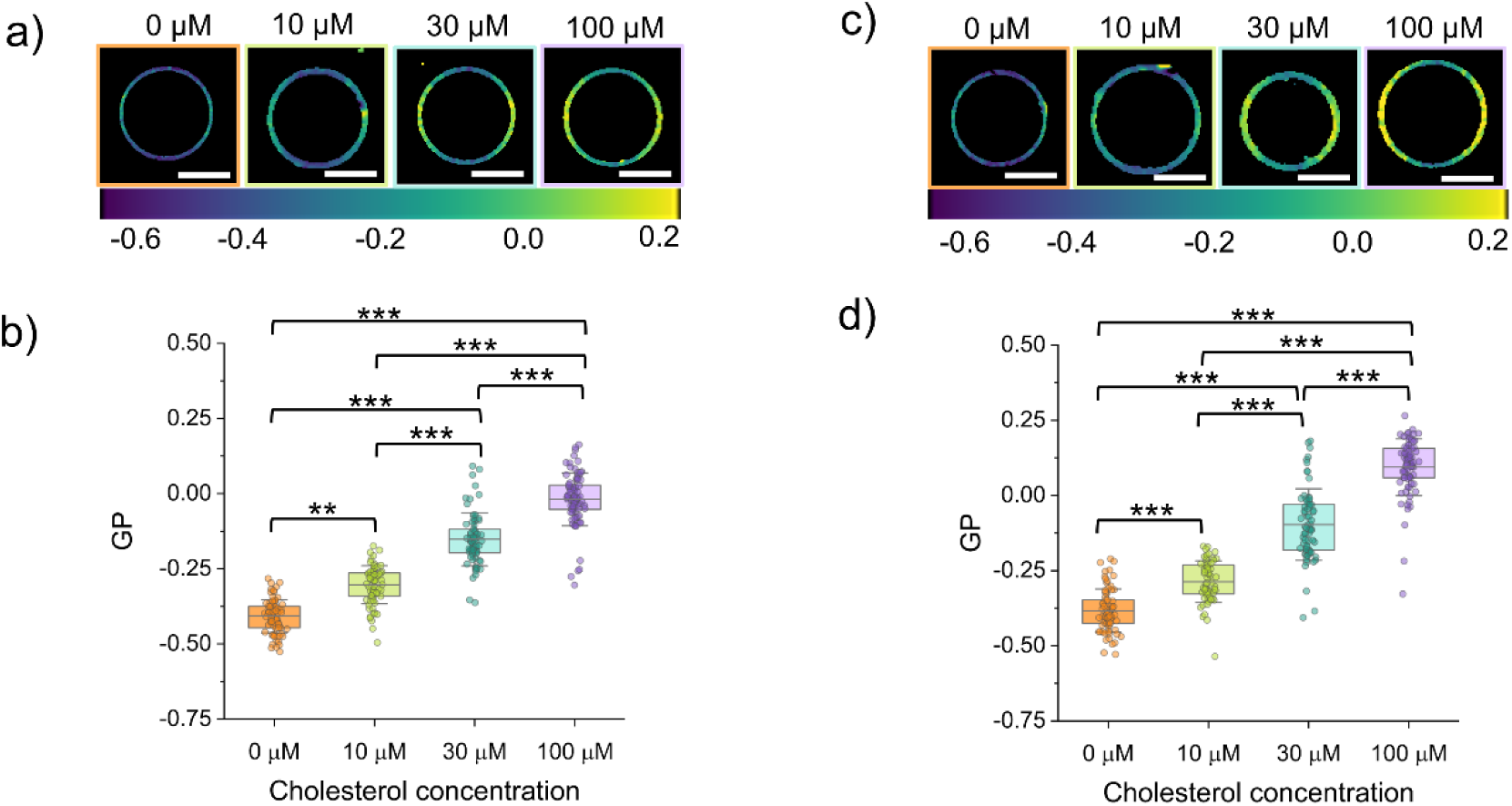
**NR12A GP measurements of cholesterol incorporation for eDICE GUVs made of binary lipid mixtures**. a) Representative color-coded GP images of DOPC:PC(18:0-14:0) (60:40 mol%) GUVs at different cholesterol (MβCD-CL) concentrations in the outer medium. b) GP values of DOPC:PC(18:0-14:0) GUVs as a function of MβCD-CL concentration. c) Representative color-coded GP images of DOPC:DMPC (6:4 mol%) GUVs at different cholesterol (MβCD-CL) concentrations in the outer medium. d) GP values of DOPC:DMPC GUVs as a function of MβCD-CL concentration. Scale bars in panels a and c indicate 10 µm. In panels b and d, box represents percentile 5-95, line in box is the mean and whiskers the SD. Points are individual GUVs (at least 50 per condition from at least 3 different replicates). Statistical significance was tested using a non-parametric Kruskal-Wallis test (** p< 0.01 and *** p< 0.001).

**Figure 5.**
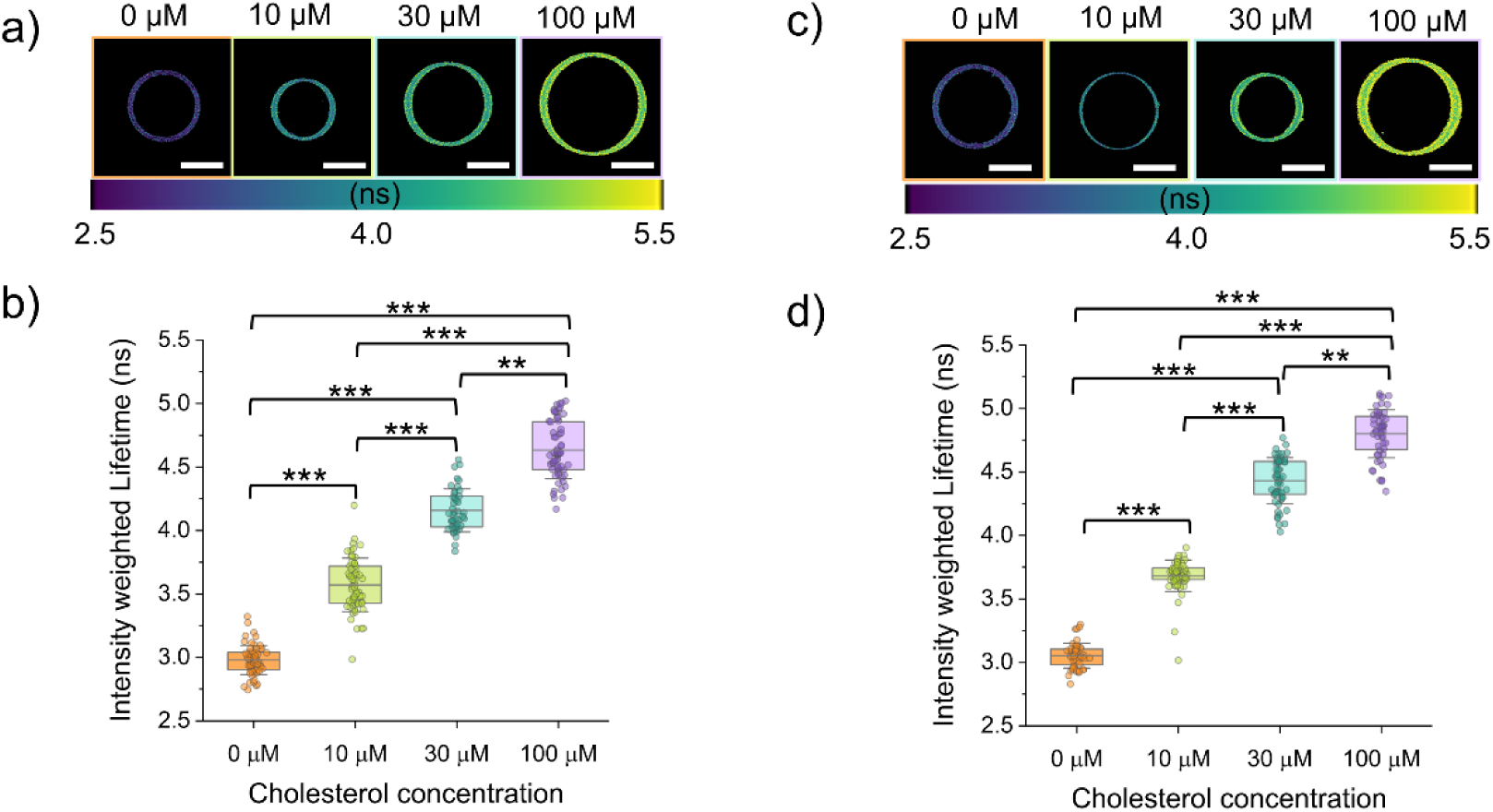
Flipper-TR fluorescence lifetimes as a reporter of cholesterol incorporation in eDICE GUVs made of binary lipid mixtures. a) Representative color- coded mean intensity weighted lifetime images of DOPC:PC(18:0-14:0) (60:40 mol%) GUVs at different cholesterol (MβCD-CL) concentrations in the outer medium. b) Mean intensity weighted lifetimes of DOPC:PC(18:0-14:0) GUVs as a function of MβCD-CL concentration. c) Representative color-coded mean intensity weighted lifetime images of DOPC:DMPC (6:4 mol%) GUVs at different cholesterol (MβCD-CL) concentrations in the outer medium. d) Mean intensity weighted lifetimes of DOPC:DMPC GUVs as a function of MβCD-CL concentration. Scale bars in panels a and c indicate 10 µm. In panels b and d, box represents percentile 5-95, line in box is the mean and whiskers the SD. Points are individual GUVs (at least 50 per condition from at least 3 different replicates). Statistical significance was tested using a non-parametric Kruskal-Wallis test (** p< 0.01 and *** p< 0.001).

Upon incubation with MβCD-CL complexes, in DOPC:PC(18:0-14:0) GUVs, the mean GP of NR12A values rose to -0.30 ± 0.06, -0.15 ± 0.09 and -0.02 ± 0.09 after treatment with 10 µM, 30 µM, and 100 µM cholesterol, respectively (Figure 4 a-b). For DOPC:DMPC GUVs, the average GP values of NR12A increased to -0.29 ± 0.07, -0.10 ± 0.12, and 0.09 ± 0.09 at the same cholesterol concentrations (Figure 4 c-d). Notably, at the highest cholesterol concentration tested (100 µM), the DOPC:DMPC mixture displayed a higher GP value than DOPC:PC(18:0-14:0), despite DMPC having a lower melting temperature. Flipper-TR lifetime measurements followed a similar trend, with the fluorescence lifetime increase upon incubation with increasing cholesterol concentrations being higher for the DOPC:DMPC GUVs (3.68 ± 0.12 ns, 4.43 ± 0.18 ns and 4.80 ± 0.19 ns after incubation with 10, 30, and 100 µM cholesterol, respectively) than for the DOPC:PC(18:0-14:0) GUVs (3.57 ± 0.21 ns, 4.15 ± 0.17 ns and 4.63 ± 0.22 ns after incubation with 10, 30, and 100 µM cholesterol, respectively) (Figure 5 a-d). The GP and fluorescence lifetime values obtained after exposure to 100 µM cholesterol are very similar to those observed in gel swollen GUVs made of DOPC:DMPC:cholesterol (36: 24: 40) (NR12A GP= 0.07 ± 0.15; Flipper-TR lifetime = 4.78 ± 0.21 ns) and DOPC:PC(18:0-14:0):cholesterol (36: 24: 40) (NR12A GP = 0.00 ± 0.07 ; Flipper-TR lifetime = 4.67± 0.20 ns) (Figure S2). These results suggest that the mismatch in chain length in PC(18:0-14:0) could reduce the efficiency of cholesterol-induced lipid packing. This is consistent with previous findings that have shown that sterols align more efficiently with symmetric saturated lipids whereas asymmetry in acyl chains disrupts this interaction, leading to a diminished effect on membrane order ^35^.

Interestingly, in our Flipper-TR experiments, we detected a small number of DOPC:DMPC GUVs exhibiting liquid-liquid phase separation after incubation with 30 µM or 100 µM cholesterol. These GUVs initially showed small bright domains with negative curvature that fused into larger and more energetically favourable domains over time (see Figure 6a and Supplementary Video 1). The final domains were clearly distinguished by their fluorescence lifetime (Figure 6b-c). Phasor analysis of the fluorescence lifetime showed that the phase separated GUVs present Lo domains with fluorescence lifetimes between 4.83 and 4.91 ns and less packed Ld domains with fluorescence lifetimes between 3.78 and 3.99 ns. Although these observations were rare (n=5 GUVs across 4 independent experiments), they suggest that cholesterol enrichment can induce phase separation in this membrane mixture. This finding highlights the potential to drive domain formation on-demand by adding cholesterol to the outer medium, which could be interesting for studying the dynamics of membrane organization in response to cholesterol or for triggering recruitment of lipid domain-specific proteins in synthetic cells ^8, 36, 37^.

**Figure 6.**
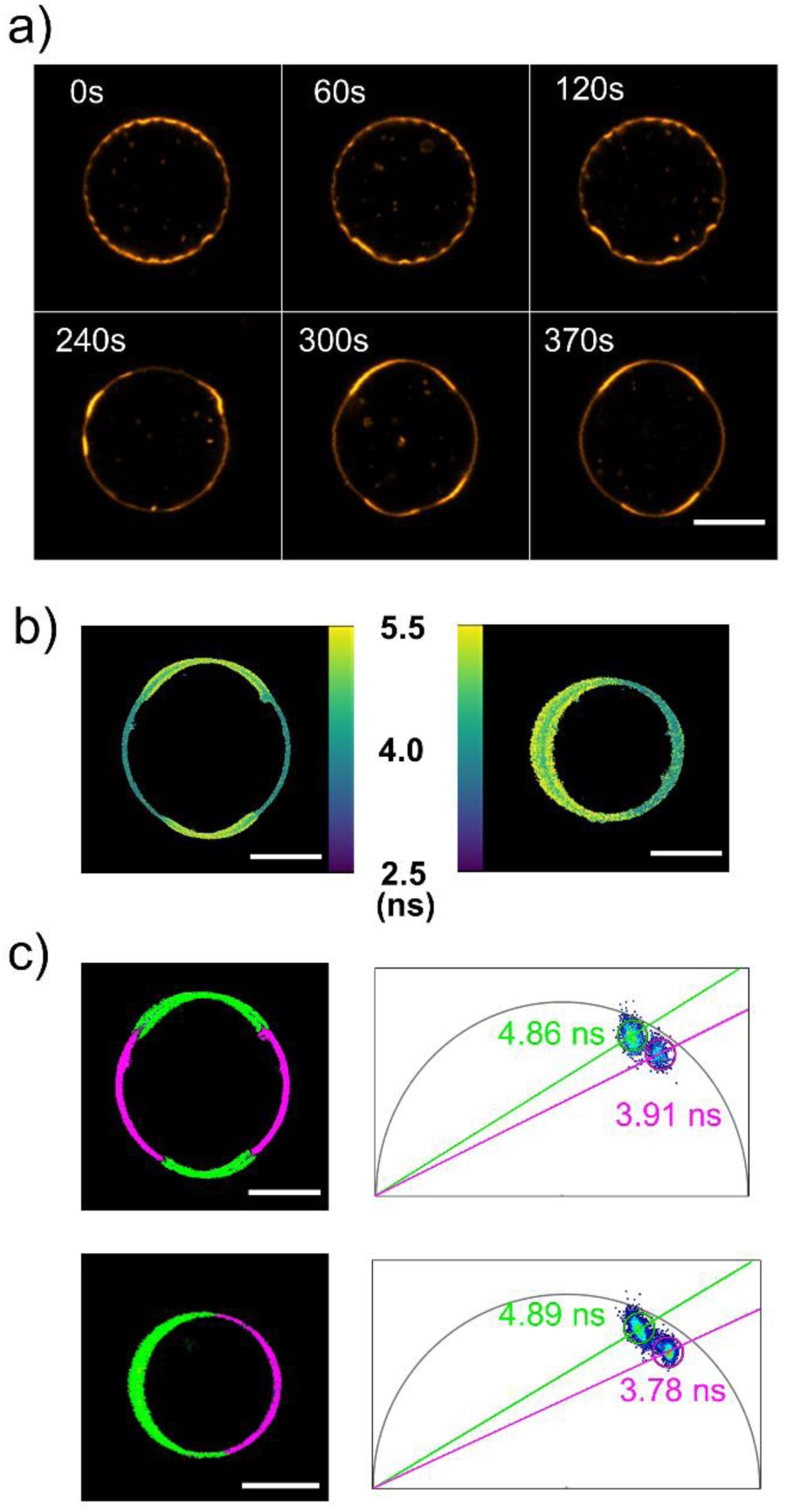
**Examples of DOPC:DMPC (6:4 mol%) GUVs exhibiting liquid-liquid phase separation upon cholesterol incorporation**. a) Confocal microscopy timelapse images of the evolution of membrane domains of a GUV labelled with Flipper-TR at different time points in the presence of 100 µM cholesterol (i.e., MβCD-CL) in the outer medium. The t=0 indicates the start of the imaging, approximately 15 minutes after the addition of MβCD-CL to the outer medium. Images show the fluorescence intensity signal of Flipper-TR at an emission wavelength between 575 and 625 nm. b) Color-coded mean intensity weighted lifetime images of two different phase separated GUVs labelled with Flipper-TR recorded within 25 and 45 minutes after addition of MβCD-CL. c) Phasor analysis of the phase separated GUVs shown in panel b. The images (left) show the regions of the membrane separated by their lifetime according to the clouds on the phasor plots on the right. The lifetime values corresponding to the Ld (magenta) and Lo (green) domains are indicated in the phasor plots. Scale bars in panels a,b,c, denote 10 µm.

## 4. Discussion

Cholesterol plays a central role in modulating the biophysical properties of cellular membranes, where it regulates membrane fluidity, packing, and domain formation. It is therefore essential to incorporate physiologically relevant cholesterol levels in model membrane systems, including giant unilamellar vesicles (GUVs), for studying membrane structure and function. Incorporating cholesterol into GUVs made by swelling is straightforward, but swelling-based methods do not allow for accurate encapsulation of cytosolic contents^6, 13^. By contrast, incorporating cholesterol into GUVs made by emulsion-based methods, which are popular for bottom-up synthetic cell reconstitution ^6, 13^, is challenging because cholesterol remains dissolved in the oil phase due to its high hydrophobicity ^10^.

The use of MβCD as a cholesterol carrier has been well established for modulating membrane cholesterol content, both in live cells^38, 39^ and in model membrane systems^7,11^. MβCD can act as both a donor and an acceptor, depending on its loading state. Our study builds on this principle by demonstrating that MβCD-CL complexes can effectively deliver cholesterol to GUVs with different initial compositions produced using the eDICE method. We demonstrated that cholesterol delivery can be reliably detected using two complementary environment-sensitive fluorescent probes: NR12A and Flipper-TR. Both probes revealed a clear, concentration-dependent incorporation of cholesterol into eDICE GUV membranes, as evidenced by increasing NR12A generalized polarization (GP) values and Flipper-TR fluorescence lifetimes. In case of DOPC GUVs, we calibrated these changes against GUVs with known cholesterol contents prepared by gel-swelling, allowing us to demonstrate tunability of the final cholesterol content of eDICE GUVs over the physiologically relevant range of 10 to 40 mol% by variations in the concentration of MβCD-CL in the outer medium. Our work shows that the NR12A and Flipper-TR probes are equally suitable for measuring the cholesterol content of GUVs. The GP values of NR12A are nonuniform across the GUV membrane due to a photoselection artifact, something that we do not observe in the Flipper-TR lifetime images. To avoid uneven fluorescence excitation of NR12A along the membrane and the resulting inhomogeneity in GP values, the microscope’s optical path can be modified by including a depolarizing component such as a quarter-wave plate, which transforms linearly polarized light into circular polarization to achieve more uniform illumination ^28^.

We observed subtle differences between the probes in GUVs composed of binary lipid mixtures with saturated lipids (DMPC and PC(18:0-14:0)). Compared to DOPC GUVs, the NR12A probe reported increased lipid packing even prior to cholesterol addition, indicating that saturated acyl chains enhance membrane order independently of sterols. However, Flipper-TR lifetimes remained similar to pure DOPC, reflecting the distinct sensing mechanisms of the two probes: NR12A responds primarily to changes in membrane polarity and local packing^16, 27^, whereas Flipper-TR detects mechanical stress and lateral tension within the bilayer^21, 22^. Upon cholesterol enrichment, both probes detected increased membrane order, with stronger responses in the DOPC:DMPC mixture compared to DOPC:PC(18:0-14:0), likely due to better cholesterol accommodation in bilayers with symmetric saturated chains^35^. Moreover, isolated observations of phase separation in DOPC:DMPC GUVs at higher cholesterol concentrations suggest that cholesterol can promote lateral heterogeneity and domain formation under specific compositional conditions. This highlights the sensitivity of membrane organization to both lipid composition and sterol content.

## 5. Conclusion

In our study we provide a unified strategy to both manipulate and monitor membrane cholesterol levels into eDICE GUVs *in situ*, by coupling post-formation cholesterol delivery via MβCD with non-invasive fluorescence microscopy using the environment-sensitive probes NR12A and Flipper-TR. This combined approach not only confirms the successful enrichment of vesicles with cholesterol but also establishes a broadly applicable method to study sterol-dependent membrane organization under controlled conditions. This method enables emulsion-based GUVs to serve as realistic, cholesterol-rich platforms for reconstitution of lipid–protein interactions and the engineering of synthetic cells.

Future studies could further investigate the emergence of phase separation by exploring a broader range of lipid mixtures, including those with higher melting temperature components such as DPPC or sphingomyelins, which are known to promote ordered domains ^7, 40^. Additionally, temperature-controlled experiments could provide critical insight into the thermodynamic conditions that favour domain formation, particularly since all current measurements were performed at room temperature. Importantly, phase-separated GUVs offer an excellent platform for reconstitution studies, where the presence of coexisting membrane domains can be exploited to control and study the spatial organization of membrane-associated proteins and cytoskeletal networks. Such systems could help unravel how lipid heterogeneity contributes to protein localization, membrane mechanics, and cellular signalling processes.

## Author contributions

MAP and GHK devised the project, conceived the main conceptual ideas and discussed the results. MAP designed and performed the experiments and data analysis and wrote the first manuscript draft. GHK contributed to the interpretation of the results, provided critical feedback and helped shape the research, analysis and manuscript.

## Data availability

Data for this article, including Flipper-TR FLIM measurements and NR12A GP values are available at 4TU.ResearchData at https://doi.org/10.4121/5a3473d9-56e7-4fe7-ba9b-cd81462ba47f The python script for the analysis of NR12A GP values of the GUVs is available on https://gitlab.tudelft.nl/marribasperez/gp-analysis-spectral-imaging.git

## Acknowledgment

We thank Robin Liu for initial experiments with the Flipper-TR probe and Nikki Nafar for helpful discussions. We acknowledge financial support from The Netherlands Organization of Scientific Research (NWO/OCW) Gravitation program Building A Synthetic Cell (BaSyC) (024.003.019) and the Kavli Synergy program of the Kavli Institute of Nanoscience Delft (to MAP).

