## Supplementary information for "Quantification of Cholesterol Incorporation in Giant Unilamellar Vesicles Produced by a Modified cDICE Method"

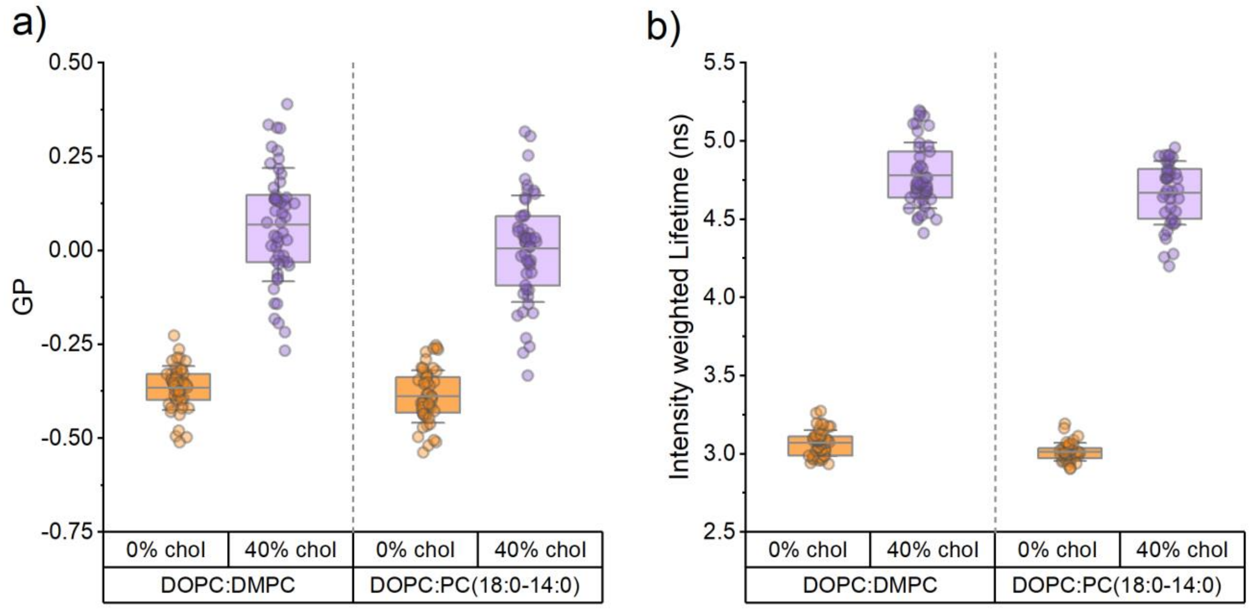


**Figure S1.** NR12A GP values (a) and Flipper-TR fluorescence lifetimes (b) of gel--swollen GUVs used for calibration. In each case, a comparison is shown between GUVs made of binary lipid mixtures without cholesterol (labelled as 0% chol) versus ternary lipid mixtures with cholesterol (labelled as 40% chol). The binary lipid compositions are DOPC:DMPC (60:40) and DOPC: PC(18:0-14:0) (60:40). The ternary lipid mixtures are DOPC:DMPC:cholesterol (36: 24: 40) and DOPC: PC(18:0-14:0): cholesterol (36: 24: 40). Box represents percentile 5-95, line in box is the mean, and whiskers the SD. Points are individual GUVs. The GUVs made of binary lipid mixtures show similar NR12A GP values and Flipper-TR fluorescence lifetimes as compared to those observed in eDICE GUVs with the same lipid composition. The NR12A GP values and Flipper-TR fluorescence lifetimes of the GUVs with a ternary lipid mixtures are comparable to the values observed in the binary eDICE GUVs after incubation with 100μM cholesterol (MβCD-CL)

**Supplementary Video 1**. Fluorescence confocal microscopy time-lapse showing phase separation of an eDICE GUV with an initial lipid composition of DOPC:DPPC (60:40 mol%) upon incubation with 100μM cholesterol (MβCD-CL) in the outer medium at room temperature. The membrane is labelled with Flipper-TR, the excitation wavelength is 488 nm and the emission is collected over a wavelength range between 575 nm and 635 nm. The recording is taken at the equatorial plane of the GUV.
